# Power-to-Vitamins: Producing Folate (Vitamin B_9_) from Renewable Electric Power and CO_2_ with a Microbial Protein System

**DOI:** 10.1101/2024.02.22.581687

**Authors:** Lisa Marie Schmitz, Nicolai Kreitli, Lisa Obermaier, Nadine Weber, Michael Rychlik, Largus T. Angenent

**Affiliations:** Environmental Biotechnology Group, Department of Geosciences, University of Tübingen, 72074 Tübingen, Germany; Chair of Analytical Food Chemistry, Technical University of Munich, 85354 Freising, Germany; AG Angenent, Max Planck Institute for Biology, Max Planck Ring 5, D-72076 Tübingen, Germany; Department of Biological and Chemical Engineering, Aarhus University, Gustav Wieds Vej 10D, 8000Aarhus C, Denmark; The Novo Nordisk Foundation CO2 Research Center (CORC), Aarhus University, Gustav Wieds Vej 10C, 8000 Aarhus C, Denmark; Cluster of Excellence – Controlling Microbes to Fight Infections, University of Tübingen, Auf der Morgenstelle 28, 72074 Tübingen, Germany

**Keywords:** Power-to-protein, CO2-to-protein, food, sustainable agriculture, yeast, microbial protein, folates

## Abstract

Meeting a surging demand for superior micronutrient-rich protein sources and finding production practices that are less detrimental to the climate will be critical challenges of the 21^st^ century. New technologies are needed to decouple food production from land use. Our group previously proposed a two-stage Power-to-Protein technology to produce microbial protein from renewable electric power and CO2. Two stages were operated *in series*: **(1)** *Clostridium ljungdahlii* in Stage A to utilize H2 to reduce CO2 into acetate; and **(2)** *Saccharomyces cerevisiae* in Stage B to utilize O2 and produce microbial protein from acetate. Renewable energy would power water electrolysis to produce H2 and O2. A disadvantage of *C. ljungdahlii* in Stage A is the need to continuously feed vitamins to sustain growth and acid production. Changing to the more robust thermophilic acetogen *Thermoanaerobacter kivui* avoids providing any vitamins. Additionally, *S. cerevisiae* produces folate when grown with acetate as a sole carbon source under aerobic conditions. A total folate concentration of 6.7 mg per 100 g biomass with an average biomass concentration of 3 g L^-1^ in Stage B is achieved. The developed Power-to-Vitamin system enables folate production from renewable power and CO2 with zero or negative net-carbon emissions.

## 1. Introduction

Current food-production practices that are based on conventional agriculture are responsible for one-quarter of global carbon emissions.[1] They rely on arable land and freshwater and are resource-extensive, leading to irreversible damage to our global environment. Land-use change by draining peatlands and deforestation decreases carbon sinks and is a considerable factor in global warming.[2] With a 30% increase in the worldwide population by 2050, the energy requirements and greenhouse gas emissions of the food sector will also rise unsustainably.[3] Simultaneously, a growing global population will exacerbate food shortages in numerous regions worldwide. The Food and Agriculture Organization of the United Nations (FAO) projections estimate that nearly 670 million people (8% of the world population) will be undernourished in 2030.[4] Countries that are already threatened by droughts and nutrient-depleted soils will be especially affected. Considering the statistics on greenhouse gas emissions from the food sector^[2a]^ and combining them with the global hunger index,[5] we can attest to an urgent requirement for a sustainable food production system. Furthermore, it will be crucial to disconnect these new food production practices from land use to preserve our ecosystems.

The brand Quorn® is a successful illustration of a microbial protein product that is currently marketed as human food. It primarily consists of mycoprotein derived from a fungal strain called *Fusarium venenatum*[6] and has gained global attention as a meat substitute for people following a vegetarian and vegan diet. Despite having a considerably lower carbon footprint than conventional meat production practices (30 times lower than beef and seven times lower than chicken),[7] mycoprotein production with glucose as a substrate still relies on agriculture for glucose production. Recently, Molitor et al. demonstrated the synthesis of microbial protein from CO2 in a two-stage bioreactor system, referred to as Power-to-Protein or CO2-to-protein, without the requirement of agriculture for protein production.[8] The two-stage bioreactor system is based on an acetate platform: **(1)** utilizing H2 to reduce CO2 into acetate by an autotrophic and anaerobic bacterium; and **(2)** oxidizing acetate with O2 into yeast biomass. The Power-to-Protein system should utilize renewable electric power to produce H2 and O2 from water splitting *via* electrolysis, which would then be fed separately into the two bioreactors *in series*.[8] The two-stage system was evaluated through thermodynamic modeling as energetically superior or competitive to other biological CO2-to-protein processes,[9] even though CO2 is lost in the acetate oxidation process in Stage B and must be recirculated to Stage A. Compared to the Quorn® production, this technology is even more sustainable, with potentially zero or even negative net-carbon emissions.[8]

In addition to implementing sustainable food production practices, considerable progress must be made in fighting micronutrient deficiencies and improving public health. Today, two billion people suffer from micronutrient deficiencies, which is defined as an insufficient supply of essential vitamins and minerals arising from poor dietary diversity.[10] The numbers are anticipated to intensify with food shortage and dilution effects on food minerals.[11] Therefore, an optimal food product should ensure balanced nutrient profiles to avoid external supplementation with vitamins and other micronutrients.

One essential vitamin of the B complex vitamins is folate (*i.e.*, vitamin B9). The structural backbone of folates consists of the pteridine molecule, which is attached to para-aminobenzoic acid, followed by a glutamate tail. Folates can be classified into different forms (vitamers) depending on: **(1)** the oxidation state of the pteridine molecule; **(2)** different Ci substituents attached at positions N^5^ and N^10^, and **(3)** the length of the glutamate tail.[12] The total folate content in a food product is always a composition of different vitamer concentrations. Common natural folates are tetrahydrofolate (H4folate), 5-methyl-tetrahydrofolate (5-CH3-H4folate), 5-formyl-tetrahydrofolate (5-CHO-H4folate), and 10-formyl-folic acid (10-CHO-H4folate).[13] In humans, folates are required for one-carbon metabolic reactions, for example, in synthesizing purine nucleotides, thymidylate (dTMP), and methionine.[14]

The active form of folate that is mainly involved in these reactions is 5-CH3-H4folate, which is the transport form in blood circulation. All folate vitamers are converted to this form during absorption through a series of enzymatic reactions. The conversion speed and efficiency of the conversion might, however, depend on the vitamer itself and the individual genetic variations in folate uptake and metabolism.[15] Regardless, any vitamer could be ultimately pertinent for human nutrition. Because humans cannot synthesize folates, they must be taken up by diet. The National Institute of Health (NIH) recommends a dietary intake (recommended daily allowance [RDA]) of 400 µg d^-1^ folate.[16] Some reasons for the insufficient supply are that folate contents in staple foods are low and the absorption of natural folate vitamers from foods is incomplete.[17] These reasons mandate a constant replenishment of folate to maintain the required levels.

In contrast to humans, many multicellular organisms, such as plants,[12] fungi,[18] and algae,[19] are capable of synthesizing folate *in vivo*. Although many microbes across the food chain can biosynthesize folates, the major staple food crops, such as rice, cassava, maize, potato, and wheat, contain insufficient folate levels.^[17a]^ Because of a high RDA of folate and an imbalanced diet, folate intake is often inadequate to maintain the required amount for humans.[20] This has inspired several strategies to counter the deficiency by pharmacological folate supplementation in the form of pills of the chemically synthesized form of folate, pteroyl-L-glutamic acid, which is also known as folic acid (PteGlu).[21] PteGlu is a popular choice for supplementation because of its stability. Unfortunately, PteGlu supplementation to an entire population is complicated to monitor and may have four limitations: **(1)** the unavailability of supplements to all relevant groups; **(2)** a lack of general awareness, and therefore an inconsistent and improper consumption; **(3)** a relatively expensive production and purification process; and **(4)** unclear adverse health effects arising from an excessive PteGlu intake.[22] Therefore, food fortification with natural folates emerged as a superior approach to improve the nutrition of entire populations.

Folate fortification entails an artificial enrichment of folates in foods either due to manual addition after the production process (*e.g.*, the addition of folates to cereal grain products),[23] breeding or genetic manipulation of plants,^[17a]^ and microbe-mediated biofortification.[24] Microbe-mediated biofortification of folate enhances the natural folate content in food by including microbes during the fermentation process.[25] Another benefit of co-production is that natural folates, compared to PteGlu, have no adverse health risks from oversaturation.[18] The direct synthesis in the food production process, moreover, avoids the extraction and purification of the folates. The baker’s yeast *Saccharomyces cerevisiae* has been reported to have high intracellular folate concentrations of approximately 1-4 mg per 100 g.[26] With a well-characterized pathway for folate synthesis,[27] *S. cerevisiae* is an ideal candidate to perform natural fortification of food by folate co-production.[28] In biotechnology, almost all yeast fermentations are based on sugar-containing substrates, including glucose, sucrose, fructose, and maltose.[29] In the Power-to-Protein system, the C2-substrate acetate is fed to yeast. To our knowledge, no reports are available that tested whether acetate as a substrate promotes vitamin (*e.g.*, folate) production in yeast.

This study aimed to analyze the folate flux within the two-stage bioreactor system composed of *Thermoanaerobacter kivui* in Stage A and *S. cerevisiae* in Stage B. One possibility is that folates are produced and released by *T. kivui* in Stage A and accumulated by the yeast in Stage B. Acetogens use H4folate as a carrier of reduced C1-fragments in the reductive acetyl-CoA pathway or the Wood-Ljungdahl pathway to enable CO2 reduction and metabolism.[30] Therefore, we expect a highly upregulated folate biosynthesis pathway. Another possibility is that folates are directly produced *de novo* by the yeast. Within this study, we could show that our system could be utilized as a Power-to-Vitamin system to biofortify food, which is essential for the sustainability and self-reliance of this mycoprotein production platform.

## 2. Experimental Section/Methods

### 2.1. Microbial strains, medium, and cultivation conditions

We cultivated pre-cultures of *T. kivui* LKT-1 (DSM 2030) in 100-mL injection bottles at 150 rpm and 60°C, containing 20 mL MS medium,[31] under strictly anaerobic conditions for 48 h. The MS-medium contained NaCl (0.45 g L^-1^), NaHCO3 (6.0 g L^-1^), K2HPO4 (0.17 g L^-1^), NH4Cl (0.19 g L^-1^), MgCl x 6H2O (0.08 g L^-1^), CaCl2 x 2 H2O (0.06 g L^-1^), KH2PO4 (0.23 g L^-1^), L-cysteine (0.5 g L^-1^), resazurin (1 mg L^-1^), 0.1% (vol/vol) trace element solution (Supporting Information [SI], Experimental procedures), 0.1% (vol/vol) (NH4)2Ni(SO4)2 (0.2%, w/v), 0.1% (vol/vol) FeCl2 x 4H2O (0.2%, w/v) and was adjusted to a pH of 6.4. We pressurized bottles to 1.0 bar overpressure with H2:CO2, 80:20 vol%. Finally, we inoculated pre-cultures with 1 mL from frozen stocks that were prepared from a previous gas fermentation study under autotrophic conditions (H2:CO2, 80:20 vol%). We never added yeast extract and vitamins to the growth media of *T. kivui*.

We pre-grew *S. cerevisiae* S288C (ATCC 204508) in 50 mL of yeast peptone dextrose (YPD) medium (yeast extract [10 g L^-1^], peptone [20 g L^-1^], glucose [20 g L^-1^]) from frozen stocks. Cultivation conditions of the yeast pre-culture were: 30°C; 150 rpm; and an overnight growth period. We performed flask experiment cultivations in 50 mL of yeast nitrogenous base medium (YNB) in 500 mL flasks with baffles. YNB consisted of (NH4)2SO4 (5 g L^-1^), KH2PO4 (1 g L^-1^), MgSO4 x 7H2O (1.025 g L^-1^), NaCl (0.1 g L^-1^), CaCl2 (0.132 g L^-1^), 10 mL trace elements (100 x), 10 mL vitamins (100 x) (SI, Experimental procedures). We supplied MES (19.52 g L^-1^) as a buffer and acetate as a substrate (200 mM, 99.7% purity) at a pH of 5.5. The flasks were inoculated with the pre-culture in YPD medium to a start OD600 of 0.05. We performed the cultivation at 30°C and 100 rpm for 96 h. During the experimental period, we took daily to determine OD600, pH, and acetate concentrations. Cultivation conditions of *C. ljungdahlii* can be found in the Experimental procedures (SI).

### 2.2 Two-stage bioprocessing system

*T. kivui* was grown in Stage A, and *S. cerevisiae* was grown in Stage B (**Figure 1**). Stage A consisted of a 2.5-L bioreactor (New Brunswick Scientific, Edison, NJ, USA) and was operated with a 2-L working volume at 63°C, a pH of 6.4, and a stirring rate at 400-800 rpm. The pH was controlled at 6.4 by an internal pH controller (Mettler-Toledo, Gießen, Germany) and regulated by adding KOH (10 M). As an additional sulfur source, a separate and continuous Na2S feed (1.54 M) was incorporated during continuous feeding with a feeding rate of 0.04 mL h^-1^. The H2/CO2 (80:20%vol) gas flow rate ranged between 100 mL min^-1^ to 200 mL min^-1^. *T. kivui* was cultivated in batch operating mode for 4 days (Period I). In continuous operation mode (Period II–VI), the reactor was fed with different concentrations of MS medium (no yeast extract and no vitamins), with a feeding rate varying between ∼40-90 mL h^-1^, corresponding to a dilution rate between 0.02 and 0.04 h^-1^ (**Table 1)**.

**Figure 1.**
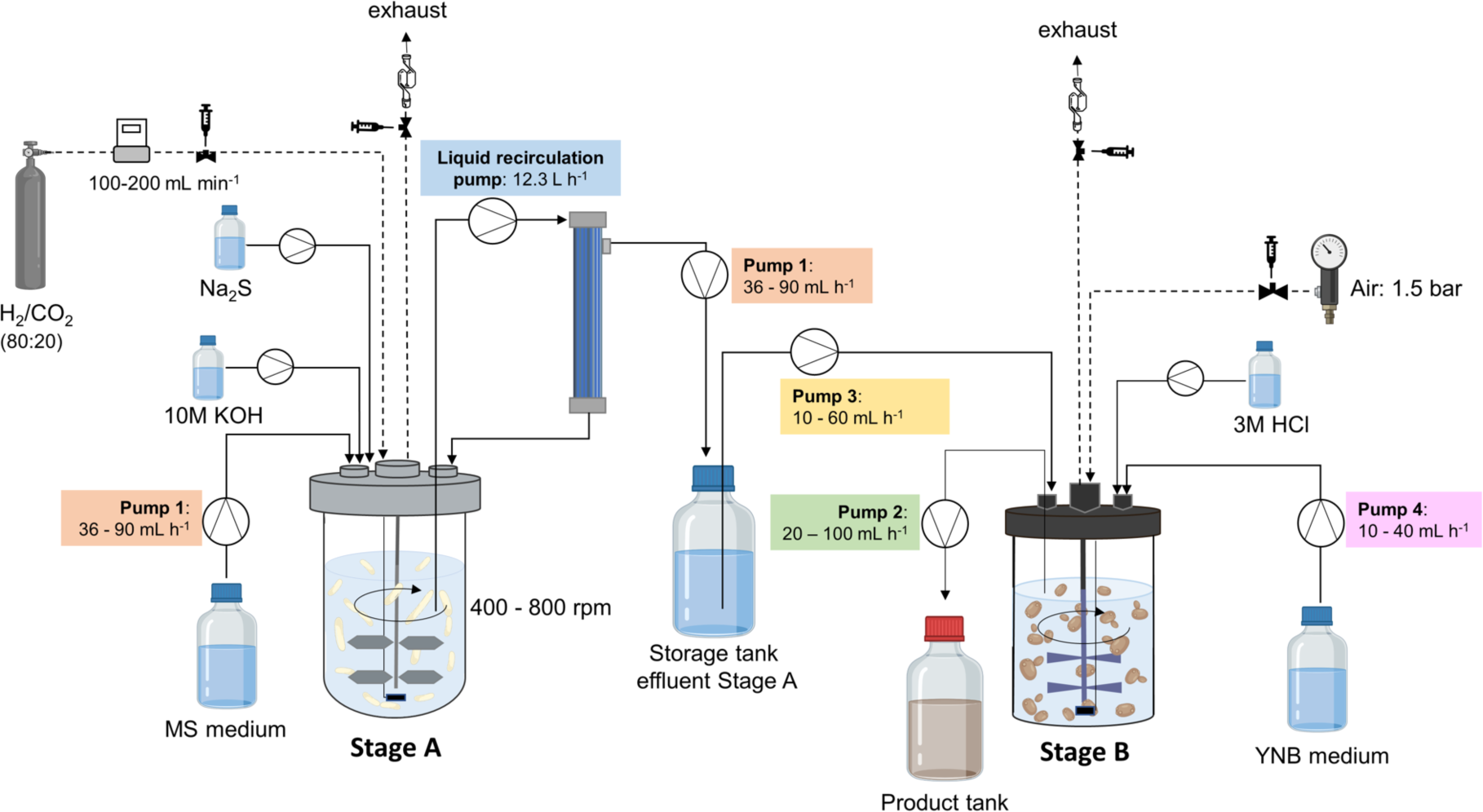
Setup of the two-stage bioreactor system. *T. kivui* was cultivated in Stage A and *S. cerevisiae* in Stage B in a continuous operating mode (figure created with BioRender).

**Table 1.** Operating parameters of the Stage A bioreactor.

| Period | Days | Operating mode | Medium concentration<sup>a)</sup> | Gas flow rate [mL min<sup>-1</sup>] | Feed rate [mL h<sup>-1</sup>] | Dilution rate [h<sup>-1</sup>] |
| --- | --- | --- | --- | --- | --- | --- |
| I | 1-4 | Batch | 1x MS | 100 | N/A | N/A |
| II | 5-14 | Continuous | 2x MS | 100 | 32 | 0.02 |
| III | 15-37 | Continuous | 2x MS | 150 | 49 | 0.03 |
| IV | 38-59 | Continuous | 4x MS | 100 | 75 | 0.04 |
| V | 60-105 | Continuous | 4x MS | 100 | 37 | 0.02 |
| VI | 106-134 | Continuous | 4x MS | 200 | 41 | 0.02 |

<sup>a)</sup> 1x, 2x, and 4x corresponds to 1-times, 2-times, and 4-times concentrated

A hollow-fiber cell recycling module (Minikros 20 cm 0.2 µM PES 1 mm 1.5TC X 3/4TC, Repligen, Breda, Netherlands) was installed to retain the bacterial cells within Stage A and to filter the effluent for Stage B. The liquid culture was circulated through the membrane with a flow rate of 12.3 L h^-1^, ensuring complete circulation of the total volume every 10 min. The filtered effluent was collected in a sterile 5-L bottle (storage tank) to decouple the two reactors and operate them at different dilution rates. We installed a temperature-dependent optical O2 sensor (PreSens Precision Sensing GmbH, Regensburg, Germany) to monitor the anaerobic status and detect unintended oxygen leaks in the reactor. The sensor was calibrated by a two-point calibration (sparging the medium with compressed air and 100% nitrogen gas).

For Stage B, we used a 2-L vessel (MyFerm, Landgraf Laborsysteme HLL GmbH, Langenhagen, Germany). We operated it with a working volume of 1.5 L at 30°C, a pH of 5.5, and a stirring speed of ∼1,000 rpm (Stuart US151 Magnetic Stirrer, Cole Parmer, Vernon Hills, IL, USA). Stage B was continuously sparged with air at 1.5 bar, which was adjusted according to the growth rate of *S. cerevisiae*. The pH was regulated at 5.5 by adding HCl (3 M). The reactor was fed with sterile effluent from Stage A (storage tank) and a YNB-bypass medium in varying concentrations in continuous operating mode. Certain specific vitamins were added as a mixture, which was not fully optimized for yeast, to the bypass flow to Stage B. Still, no yeast extract, no folate, and no cobalamin (*i.e.*, vitamin B12) were present. Stage A effluent and medium ratios were applied (**Table 2)**. We continuously connected the product in a 5-L brown glass bottle. The dilution rate for Stage B varied between 0.02 and 0.07 h^-1^.

**Table 2.** Feeding ratios of Stage A effluent and bypass medium throughout the cultivation period of 134 days.

| Period | Operating mode | Days | Stage A-effluent<br>feed [%] | Medium [%] |
| --- | --- | --- | --- | --- |
| i | Batch | 6-9 | 50 | 50 (2xYNB<sup>a)</sup>) |
| ii | Continuous | 10-22 | 50 | 50 (2xYNB) |
| iii | Continuous | 23-35 | 70 | 30 (2xYNB) |
| iv | Continuous | 36-55 | 70 | 30 (4xYNB<sup>b)</sup>) |
| v | Continuous | 58-136 | 60 | 40 (4xYNB) |

a) 2-times concentrated, b) 4-times concentrated

### 2.3. Analytical procedure

Samples from Stage A and Stage B were collected daily, and the growth (OD600), substrate consumption (GC), and product formation (HPLC) were analyzed. We monitored the change in biomass concentration (batch and continuous cultivation) by optical density (OD600) measurements at 600 nm (BioMate™ 160 UV-Vis Spectrophotometer, Thermo Fisher Scientific, Waltham, MA, USA). Cell dry weight (CDW) was determined by using correlation factors of 354.9 mg CDW OD600^-1^ L^-1^ for *T. kivui* and 515 mg CDW OD600^-1^ L^-1^ for *S. cerevisiae,* as defined in this study. Samples from both stages were monitored daily with a phase contrast microscope (Olympus BX41, Olympus, Tokyo, Japan) to check contaminations in the bioreactors. In addition, we checked the pH in both bioreactor broths with an external pH probe after taking a daily sample (VWR pHenomenal® pH 1100L, Radnor, PA, USA). Acetate concentrations were measured using high-pressure liquid chromatography (HPLC) for both bioreactors (LC20, Shimadzu, Kyoto, Japan) with an Aminex HPX-87H column (Bio-Rad, Hercules, CA, USA) as a stationary phase. 5 mM sulfuric acid was used as a liquid phase. The method was run isocratic for 31 min, and the injection volume of the samples was 20 µL. The term acetate in this paper always refers to a mixture of acetate and acetic acid because it is pH-dependent.

Gas samples were collected at the inlet and the outlet of the gas lines to analyze the relative partial pressures of the gases (H2/CO2 for Stage A; N2/O2/CO2 for Stage B) by GC as described in the supporting information (SI, Experimental procedures). Inlet gas flow rates were measured by mass flow controllers (200 SCCM, Alicat Scientific, Tucson, AZ, USA). Outlet gas flow rates were measured by a drum-type gas meter (TG 0.5, Ritter, Bochum, Germany). The gas pressure in the headspace of the bioreactors was determined by a digital pressure gauge (Cole Parmer, Vernon Hills, IL, USA). The partial pressures, the volumetric flow rates, and the gas pressure were used to determine the gas consumption and production rates. Ammonium concentrations in the medium and effluent bottle of Stage A were analyzed by the Nessler method, and trace element concentrations were measured by inductively coupled plasma mass spectrometry (ICP-MS) (SI, Analytical methods).

### 2.4. Protein determination

We took 2-mL samples for protein determination. The samples were centrifuged at 13,000 rpm, washed with 1 mL of ddH2O, and dissolved in 1 mL of NaOH (1 M). The samples were stored at -20°C until the protein measurement. Protein concentrations were measured by the BCA protein assay[32] using the Pierce™ BCA Protein Assay Kit (Thermo Fisher Scientific™, Waltham, MA, USA) and by protein determination according to Lowry[33] (SI, Experimental procedures).

### 2.5. Folate analysis

Biomass and supernatant samples were collected from Stage A and Stage B. Samples from Stage A (*T. kivui*) were taken on day 134 by harvesting the whole reactor broth. Stage B (*S. cerevisiae*) samples were taken from the product tank after collecting effluent for two days on days 94 and 126. Samples were centrifuged at room temperature and 3,430 g for 10 min. The supernatant was transferred to a new tube. All supernatant and pellet samples were immediately stored at -80°C before freeze-drying (biomass samples) (SI) or until analysis (liquid samples).

All samples were analyzed *via* stable isotope dilution assays (SIDA) after Striegel et al. [34] and Obermaier et al.[35] All buffers and solutions can be found in the previously mentioned publications. 10 mg of biomass and 500 µL of supernatant samples were weighed for analysis and mixed with extraction buffer. ^13^C-isotope-labeled internal standards (ITSD) were added to all the samples for five folate vitamers: **(1)** [^13^C5]-PteGlu; **(2)** [^13^C5]-H4folate; **(3)** [^13^C5]-5-CH3-H4folate; **(4)** [^13^C5]-5-CHO-H4folate; and **(5)** [^13^C5]-10-CHO-PteGlu. The amount of ITSD was added based on the expected folate concentration within each sample. After equilibration, an enzymatic deconjugation of the folate polyglutamates to its monoglutamic form was performed using chicken pancreas and rat serum overnight.[34–35] Afterward, the folates were further purified through solid-phase extraction with an anion-exchange column (Strata SAX cartridges, quaternary amine, 500 mg, 3 mL, Phenomenex Aschaffenburg, Germany) and eluted with 4 mL elution buffer. The eluent was filtered (PVDF, 0.22 µm), and measured on a triple quad LC-MS/MS system (Shimadzu Nexera X2 UHPLC system, Kyoto, Japan) using an ARC-18 column (2.7 μm, 100 mm × 2.1 mm, Restek, Bad Homburg, Germany).

## 3. Results and Discussion

### 3.1. *T. kivui* achieved industrially relevant acetate production rates without supplying any yeast extract and vitamins

A previous proof-of-concept study by Molitor et al. showed that *C. ljungdahlii* in Stage A achieved a maximum volumetric acetate production rate of 1 g L^-1^ h^-1^ during an operating period of 104 days.[8] However, at the beginning of the fermentation period, the P7 medium included yeast extract supplementation. In addition, a complex vitamin solution containing folate and cobalamin was fed throughout the entire fermentation period to promote bacterial growth.[8] To run the system sustainably, we envision providing only the indispensable vitamins and nutrients, allowing the microbes to produce the other vitamins themselves. Optimally, no yeast extract and no vitamins would need to be added. Here, we were especially interested in the enrichment of folate. Unfortunately, we could not grow *C. ljungdahlii* in the Stage A bioreactor without yeast extract and the vitamins folate and cobalamin in the P7 medium. After an adaptation period in batch (I) and early continuous mode (II), we still fed P7 medium containing yeast extract (1 g L^-1^ and 0.5 g L^-1^, respectively) and the vitamins folate and cobalamin. After omitting yeast extract and folate from the medium (III), we observed a decrease in the biomass concentration, a drop in the acetate production rate, and an impediment in the consumption of the gases H2 and CO2 (data shown in SI, **Figure S1**). Folate is a cofactor that was predicted to be synthesized from genomic information of *C. ljungdahlii*.[36] However, in our hands, growth without folate was cumbersome for *C. ljungdahlii*. Others recently reported that *C. ljungdahlii* can grow in a mineral medium without yeast extract after the omission of folic acid. However, the authors added vitamin-free casein acid hydrolysate, which may still provide precursors for vitamin synthesis.[37] Thus, *C. ljungdahlii* in Stage A is not an ideal acetogenic bacterium due to the requirement for yeast extract, folate, or cobalamin.

*T. kivui* is a thermophilic acetogenic bacterium that was initially isolated from Lake Kivu between the Democratic Republic of the Congo and Rwanda.[38] The advantage of *T. kivui* is that the strain has short doubling times of 1.75-2.5 h during growth on H2 + CO2, while it does not require any vitamins.[38] As the prototrophy for vitamins corresponded to our demands, we cultivated *T. kivui* in the Stage A bioreactor with *S. cerevisiae* in the Stage B bioreactor in series. Because *T. kivui* is a thermophilic microbe, we operated the Stage A bioreactor at 63°C. After a batch period of four days (Period I), the reactor was operated continuously (Period II– VI) throughout an operating period of 135 days (**Table 3**). The biomass concentration remained at approximately 0.5 gCDW L^-1^ during Periods II–IV and 1.0 gCDW L^-1^ during Periods V–VI (Figure 2A). Thus, the biomass concentration was the highest at the end of the study (Period VI in Figure 2A). Because we integrated a cell-retention system in Stage A, the growth rate was independent of the dilution rate. At this point, we did not distinguish between metabolic-active and dead cells. Therefore, increased cell concentrations could also result from an accumulation of dead cells because they were not removed from the bioreactor. However, in tandem with an increased biomass concentration, we observed a slow increase in the volumetric acetate production rate. For example, during Period VI, for which we observed the highest biomass concentration (Figure 2A), we achieved a maximum volumetric acetate production rate of 0.63 g L^-1^ h^-1^ ± 0.07 g L^-1^ h^-1^ (0.5 mol-C L^-1^ d^-1^) (Figure 2B) at an average CO2 consumption rate of 0.99 ± 0.33 g L^-1^ h^-1^ (0.54 mol-C L^-1^ d^-1^) (Figure 2C). In addition, a maximum titer of 600 mM acetate was achieved during Period VI (**Figure S2, SI**). The average base (KOH) consumption in Period VI was 0.9 g L^-1^ h^-1^. Because acetate is the sole product of the metabolism of H2 and CO2 with *T. kivui*,[38] we did not observe carbon losses to ethanol. Consequently, even though dead cells might have accumulated, a high concentration of metabolically active cells was still present.

**Figure 2.**
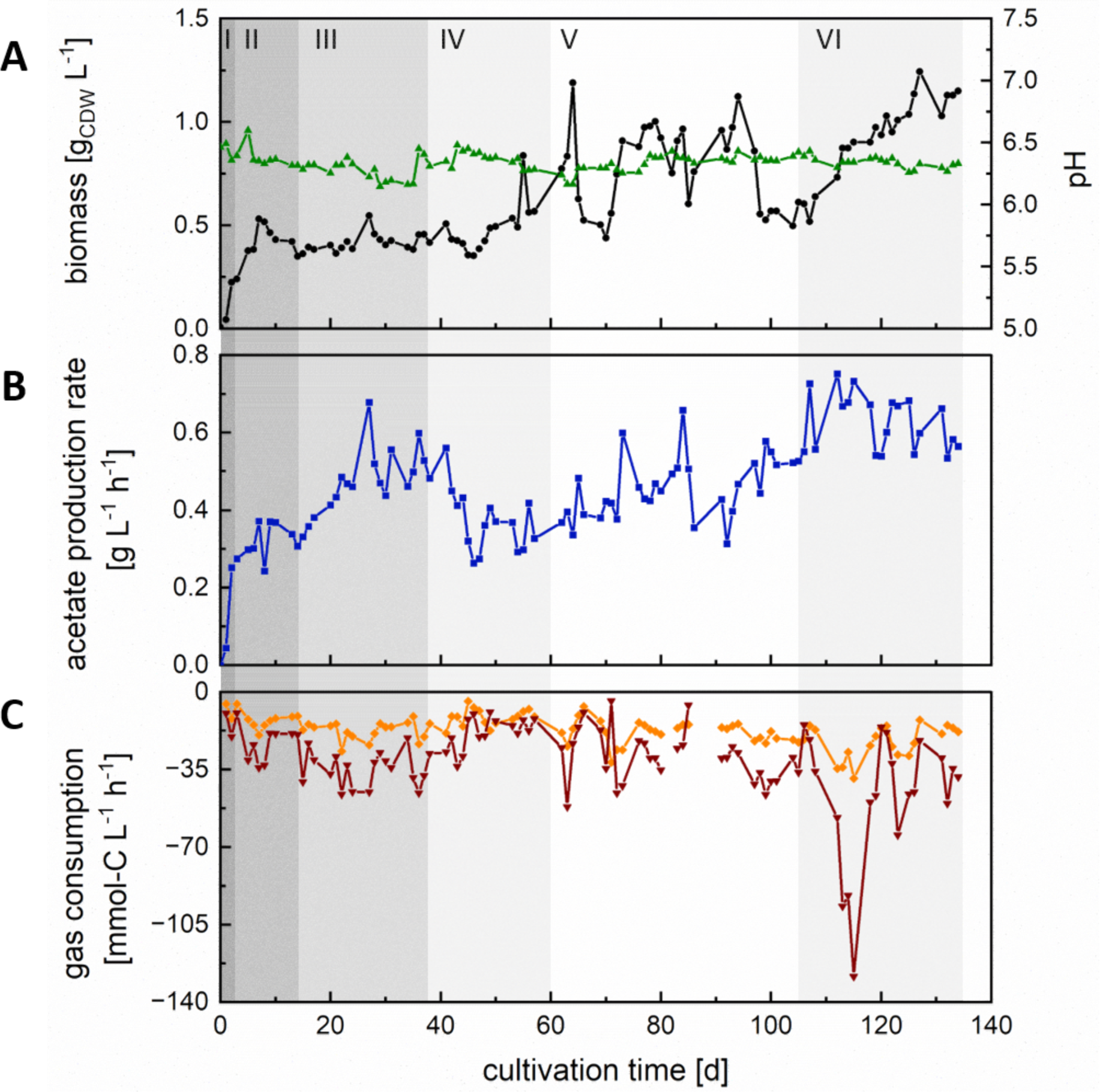
Performance of Stage A with *T. kivui* during the continuous operating period of 134 days. (A) Biomass in grams cellular dry weight (CDW) per liter (black) and pH (green); (B) Production rate of acetate in g L^-1^ h^-1^ (blue); (C) Consumption rates for H_2_ (red) and CO_2_ (orange) in mmol-C L^-1^ h^-1^. Period I: batch operation; Period II–VII: continuous operation. Period II: gas flow rate of 100 mL min^-1^ and dilution rate of 0.02 h^-1^, 2x MS medium; Period III: gas flow rate 150 mL min^-1^ and dilution rate of 0.03 h^-1^, 2x MS medium; Period IV: gas flow rate 100 mL min^-1^ and dilution rate of 0.04 h^-1^, 4x MS medium; Period V: gas flow rate 100 mL min^-1^ and dilution rate of 0.02 h^-1^, 4x MS medium; Period VI: gas flow rate 200 mL min^-1^ and dilution rate of 0.02 h^-1^, 4x MS medium.

**Table 3.**
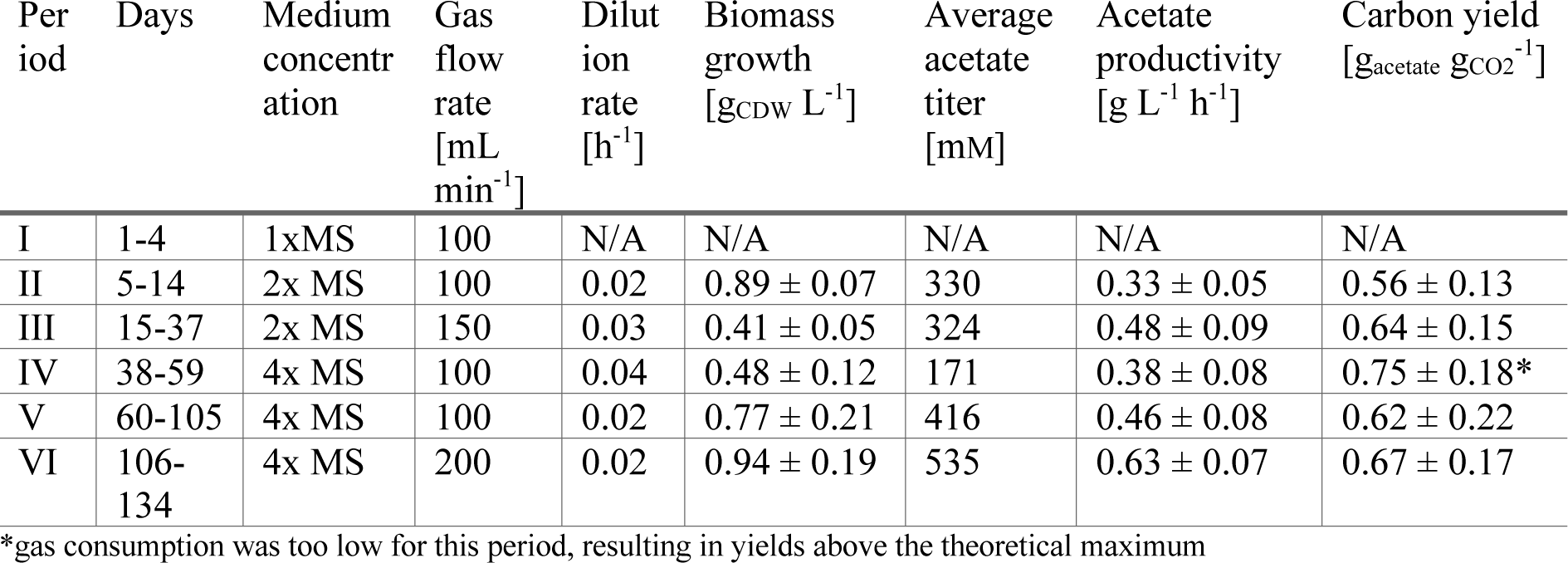
Process parameters and performance of Stage A bioreactor in the Periods I–VI.

| Per<br>iod | Days | Medium<br>concentr<br>ation | Gas<br>flow<br>rate<br>[mL<br>min<sup>-1</sup>] | Dilut<br>ion<br>rate<br>[h<sup>-1</sup>] | Biomass<br>growth<br>[g<sub>CDW</sub> L<sup>-1</sup>] | Average<br>acetate<br>titer<br>[mM] | Acetate<br>productivity<br>[g L<sup>-1</sup> h<sup>-1</sup>] | Carbon yield<br>[g<sub>acetate</sub> g<sub>CO2</sub><sup>-1</sup>] |
| --- | --- | --- | --- | --- | --- | --- | --- | --- |
| I | 1-4 | 1xMS | 100 | N/A | N/A | N/A | N/A | N/A |
| II | 5-14 | 2x MS | 100 | 0.02 | 0.89 ± 0.07 | 330 | 0.33 ± 0.05 | 0.56 ± 0.13 |
| III | 15-37 | 2x MS | 150 | 0.03 | 0.41 ± 0.05 | 324 | 0.48 ± 0.09 | 0.64 ± 0.15 |
| IV | 38-59 | 4x MS | 100 | 0.04 | 0.48 ± 0.12 | 171 | 0.38 ± 0.08 | 0.75 ± 0.18* |
| V | 60-105 | 4x MS | 100 | 0.02 | 0.77 ± 0.21 | 416 | 0.46 ± 0.08 | 0.62 ± 0.22 |
| VI | 106-134 | 4x MS | 200 | 0.02 | 0.94 ± 0.19 | 535 | 0.63 ± 0.07 | 0.67 ± 0.17 |

\*gas consumption was too low for this period, resulting in yields above the theoretical maximum

Compared to the previous proof-of-concept study by Molitor et al., the acetate production rate was only 64% of the maximum acetate production rate with *C. ljungdahlii* (0.98 g L^-1^ h^-1^ and 0.78 mol-C L^-1^ d^-1^).[8] Another study used the anaerobic thermophilic bacterium *Moorella thermoacetica* to produce acetate from H2 and CO2 as an intermediate for lipid-producing yeast.[39] The volumetric acetate production rate for *M. thermoacetica* was 0.9 g L^-1^ h^-1^, and therefore higher than in our study. Even though the authors did not supplement the medium with individual vitamins, the medium contained 10 g L^-1^ yeast extract, which includes a complex and undefined mixture of vitamins. However, the MS medium the we used in our study only contained mineral and trace salts, ammonium chloride as a nitrogen source, and cysteine and sodium sulfide as a sulfur source. We did not add yeast extract or vitamins to the medium for Stage A. Our values are comparable, but slightly lower, to an acetate production rate of 0.76 g L^-1^ h^-1^ with *T. kivui*, which was recently published.[40] These authors reported a maximum acetate titer of 490 mM and also did not apply yeast extract or vitamins.

Increasing the dilution rate (Period IV compared to Period II) and doubling the medium concentration from 2-fold to 4-fold (Period V compared to Period II) only had a minor effect on cell growth and acetate productivity for Stage A (**Table 3**). By analysis of ammonium concentrations and trace element concentrations in the medium and effluent of Stage A, we showed that nutrients were never limited (SI, **Table S1**). This verifies that an increased medium supply cannot support higher growth. Because the highest volumetric acetate production rate was observed when the gas flow with 200 mL min^-1^ was the highest (Period VI), we anticipate that gas transfer was the major limitation of the volumetric acetate production rate. Moreover, the carbon yield from CO2 to acetate for Stage A had an average value of 65 ± 20% (n = 92), which meets the theoretical maximum of 66% considering the stoichiometry according to Mishra *et al*.[9] This confirms that the process was not limited by nutrient availability but was dependent on dissolved gas availability. A considerable limitation of gas fermentations is the mass transfer of the gas H2 into the liquid.[41] We, therefore, expect a substantial increase in the production rates if we overcome mass transfer limitations.

### 3.2. *T. kivui* enriches high amounts of folate inside the cells, but folate is not released in the medium

In the methyl branch of the Wood-Ljungdahl pathway, CO2 is reduced to formate and attached to a one-carbon (C1) carrier. In acetogens, H4folate serves as the primary carrier of C1 units.[42] The formate as a C1 unit is loaded to the N10 position of H4folate (catalyzed by the formyl-H4folate synthetase). This ATP-requiring step of the Wood-Ljungdahl pathway leads to 10-CHO-H4folate, which is further reduced to 5,10-CH^+^-H4folate, 5,10-CH2-H4folate (by a formyl-H4folate cyclohydrolase/methylene-H4folate dehydrogenase), and finally, 5-CH3-H4folate (by a methylene-H4folate reductase). Accordingly, a high folate content in *T. kivui* is not surprising. Analysis of the total folate content in our *T. kivui* biomass revealed a total folate concentration of 13 mg per 100 g biomass (Figure 3**; Table S2, SI**), which is at a minimum eight times higher than the intracellular concentration of other described folate-producing wild-type bacteria.[43] The most abundant vitamers were H4folate (25 %), PteGlu (27 %), 5-CH3-H4folate (27 %), and 5-CHO-H4folate (19 %), accounting for approximately 98% of the total folate content (Figure 3). The PteGlu vitamers (PteGlu and 10-CHO-PteGlu) detected in the samples are the fully oxidized forms of the corresponding reduced H4folate forms (H4folate and 10-CHO-H4folate) and presumably resulted from an interconversion reaction occurring during sampling or sample preparation due to pH adjustments. PteGlu has no bioactive function, and its presence does not appear to benefit the microbe. A detailed discussion on the vitamer extraction process and observed interconversion reactions can be found in Gmelch *et* al.[44]

**Figure 3.**
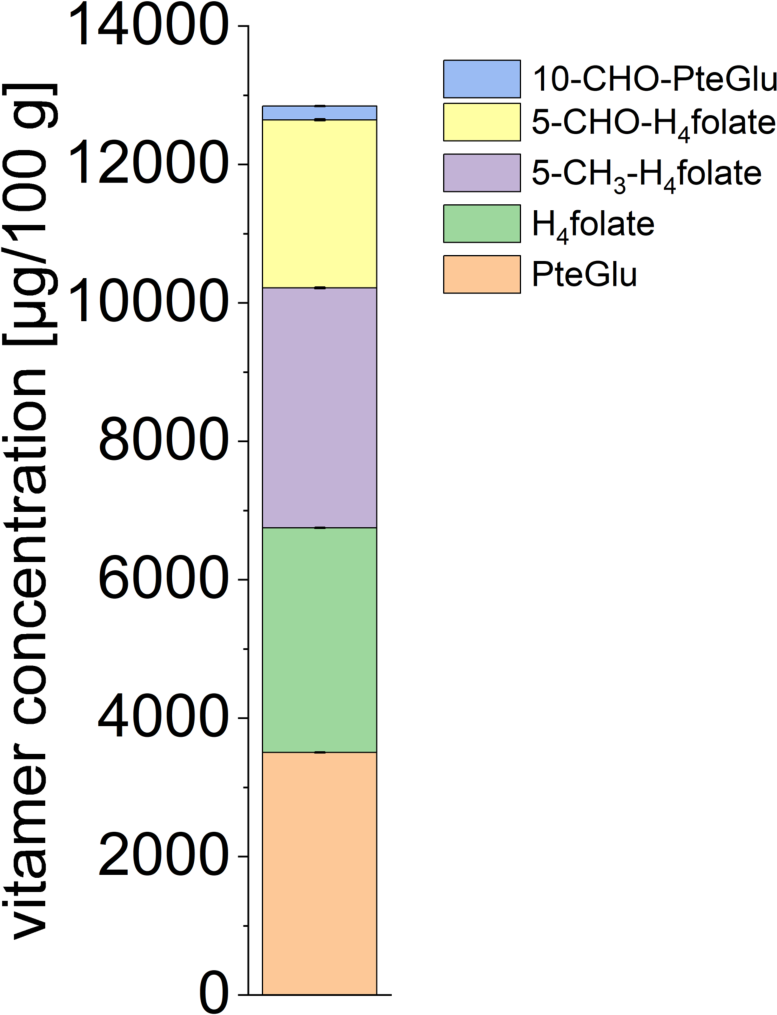
Total folate contents in biomass of *T. kivui*. For the vitamin analysis, each extraction was conducted in duplicates. The vitamers PteGlu, H_4_folate, 5-CH_3_-H_4_folate, 5-CHO-H_4_folate, and 10-CHO-PteGlu were measured. Concentrations are given in µg 100 g_CDW-1_ with standard deviations in %.

While the relatively high fraction of H4folate and 5-CH3-H4folate can be explained with them being the primary intermediates of the Wood-Ljungdahl pathway,[42, 45] 5-CHO-H4folate is not a standard intermediate of the Wood-Ljungdahl pathway. As described above, normally, the C1 unit is transferred to the N10 position of H4folate, resulting in 10-CHO-H4folate, which, however, in our analysis, was only present at about 1.5% (measured as 10-CHO-PteGlu in our study) of the total folate content. Some C1 units can be transferred to the N5 position of the H4folate carrier molecule, leading to 5-CHO-H4folate.[42] Yet, 10-CHO-H4folate is the more favorable substrate of the formyl-H4folate cyclohydrolase,[42] because reducing 5-CHO-H4folate compared to 10-CHO-H4folate to 5,10-CH^+^-H4folate requires an additional ATP.[46] Consequently, a faster conversion of 10-CHO-H4folate would explain the lower detected levels of 10-CHO-H4folate and an accumulation of 5-CHO-H4folate.

In contrast to high intracellular folate levels, the folate concentration in the medium was only ∼ 3.2 µg L^-1^, indicating that almost all the folate is stored inside the cells of *T. kivui* and is not released to the medium. To further enrich the folate titers of *S. cerevisiae*, we explored if we can feed cell supernatant from disrupted *T. kivui* cells, employing both sonication and bead mill methods. Unfortunately, we were not able to extract the vitamers from *T. kivui* to the cell supernatant. Furthermore, there is no evidence that *S. cerevisiae* can take up folates from the environment,[47] even though PteGlu is usually added as a component in minimal media for yeasts. To use the high folate levels in *T. kivui* for food fortification, an option could be to add a small amount of *T. kivui* cells to the final protein product from *S. cerevisiae*. Another option would be to separate folate from *T. kivui* through industrial methods and to add extracted folate to the final product. However, further investigations are needed.

### 3.3. *S. cerevisiae* produced folates when fed acetate under aerobic conditions

Although it is already known that *S. cerevisiae* can synthesize folates when fed with glucose,[48] one of the scopes of this study was to determine whether this yeast can produce folates when grown with acetate as a carbon source. Sugar phosphates are the starting substrates for the purine nucleotide synthesis pathway, involving the pentose phosphate pathway to produce the pterin moiety of folates. Thus, with glucose as a substrate, the sugar moiety is already present. However, with acetate as a substrate, an anabolic pathway for carbon assimilation is necessary. This may affect the carbon flow into folate synthesis, possibly yielding a lower folate content in yeast with acetate than with glucose. Acetate is first converted by an acetyl coenzyme A synthetase (ACS) to acetyl-CoA and can be either directly oxidatively metabolized in the TCA cycle in the mitochondrion or can enter the glyoxylate cycle and the anabolic gluconeogenesis pathway to form sugar phosphates.[49] Therefore, we wanted to investigate whether folate synthesis is possible while feeding only acetate as a carbon source (SI, **Figure S3**). We performed batch-cultivation and continuous-reactor experiments. First, for batch-cultivation experiments, folate production was evident in *S. cerevisiae* when pure acetate as a sole carbon source was provided in YNB medium (b_f1, b_f2 in Figure 4). We observed concentrations of ∼3.3 mg per 100 g biomass. H4folate and 5-CH3-H4folate constituted almost 90% of the total folate content, with 35% and 53%, respectively. This distribution is consistent with other studies in which 5-CH3-H4folate and H4folate were the main vitamers after batch cultivation with glucose as a substrate, with up to 80% 5-CH3-H4folate of the total folate content.[28, 48, 50] To the best of our knowledge, this is the first study demonstrating folate biosynthesis from yeasts that was cultivated on acetate.

**Figure 4.**
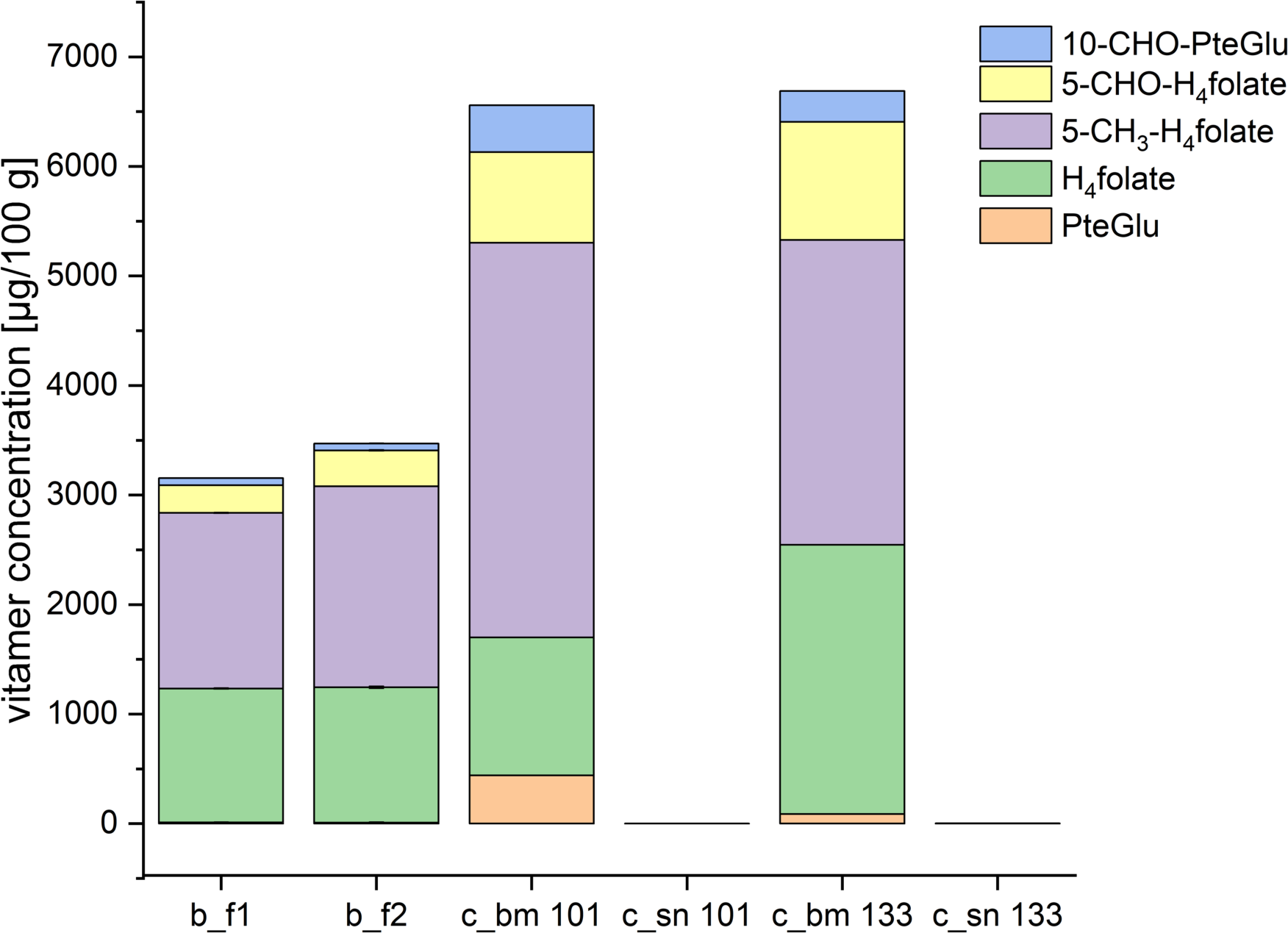
Total folate contents in biomass and medium as a sum of different vitamers. Samples were taken from flask 1 and 2 from batch-cultivation experiments (b_f1 and b_f2), and the continuous-reactor experiment after Day 101 and Day 133 (c_101 and c_133). Samples were taken from biomass (bm) and the supernatant (sn). For the vitamin analysis, each extraction was conducted in duplicates. The vitamers PteGlu, H_4_folate, 5-CH_3_-H_4_folate, 5-CHO-H_4_folate, and 10-CHO-PteGlu were measured. Concentrations are given in µg 100 g_CDW-1_ with standard deviations in %.

Second, we tested the yeast strain in the continuous-reactor setup, coupling the Stage B bioreactor to the Stage A bioreactor. We hypothesized that growth and intracellular folate accumulation by *S. cerevisiae* is possible with acetate that was produced in Stage A by *T. kivui*. At the beginning of the cultivation, we observed an incomplete acetate consumption. An oxygen limitation most likely caused the insufficient consumption of acetate due to clogging of the sparging stone after a three-week operating period (Day 28, marked by arrow). After replacing the sparging stone and restarting the reactor on Day 95, a 100% acetate consumption and a maximal biomass production rate of 0.24 gCDW L^-1^ h^-1^ (Figure 5A) were reached. The acid consumption (HCl) at 100% acetate consumption was approximately 0.56 g L^-1^ h^-1^. The protein mass fraction determined by the Lowry method was approximately 40% (Figure 5C), which is consistent with values reported in other studies.[51] This yielded a maximum protein production rate of 0.1 gprotein L^-1^ h^-1^. The biomass and protein production rates for our study were slightly higher than for the previous study by Molitor et al., who reported biomass and protein production rates of 0.18 gCDW L^-1^ h^-1^ and 0.07 gprotein L^-1^ h^-1^, respectively.[8]

**Figure 5.**
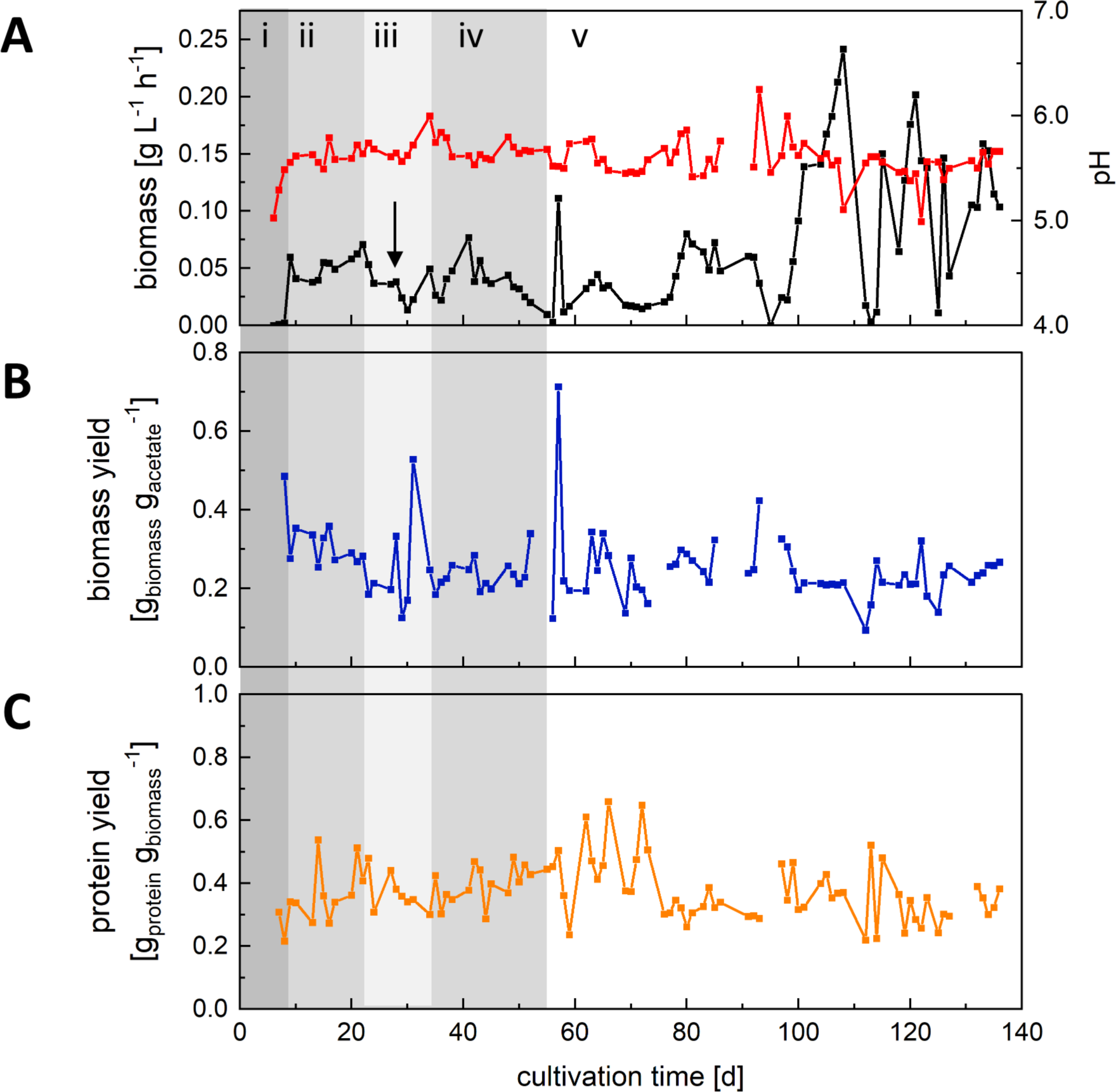
Performance of Stage B with *S. cerevisiae* during the continuous operating period of 136 days. (A) The biomass production rate in g L^-1^ h^-1^ (black) and pH (red), (B) biomass yield in g_biomass_ g_acetate-1,_ and (C) protein yield g_protein_ g_biomass-1_. The arrow marks clogging of the sparging stone.

Unfortunately, we always observed a wash-out of yeast cells at a dilution rate beyond 0.06 h^-1^. This had already been observed before,[8] even though the maximum growth rate for *S. cerevisiae* on acetate *via* the glyoxylate cycle was reported to be 0.31 h^-1^.[52] With a balanced growth rate (µ) and dilution rate (D) in chemostat conditions, theoretically, there is potential to enhance dilution rates further. A limited oxygen supply might still have hindered high biomass production rates for us and needs further investigation. In the future, the two-stage system will be coupled to water electrolysis, and the supply of pure oxygen could overcome oxygen limitations. Additionally, the medium supply could be further improved. In the described setup, we added YNB medium as a bypass medium containing vitamins to enhance the growth of *S. cerevisiae*. Some literature suggests that the *S. cerevisiae* pangenome harbors all the necessary genetic information to synthesize most of the vitamins itself. However, the growth rate can be increased by an additional supply of vitamins, thereby, overcoming the metabolic burden of synthesizing the vitamins.[53] In further experiments, the necessary supply of vitamins should be investigated in detail to minimize the use of procured vitamins and maximize growth. On average, the achieved biomass yield (gbiomass gacetate^-1^) of 0.28 g g^-1^ (Figure 5B) was consistent with Verduyn et al., who determined a biomass yield of 0.29 g g^-1^ for *S. cerevisiae* on acetate.[54]

Several parameters determine the folate composition in *S. cerevisiae*, such as the medium composition, nutrient availability, and growth conditions (*e.g.*, batch *vs*. continuous growth). On the one hand, Hjortmo et al. observed a higher cellular folate content after growth in a synthetic medium; on the other hand, they determined folate concentrations to be growth rate-dependent.[28] Higher folate contents were reached in fast-growing cells compared to slow-growing cells. Batch cultivation has the inherent disadvantage that the highest growth is only temporary when high substrate concentrations are available. On the contrary, chemostats can be operated at steady-state conditions and can control the growth rate close to the maximum. Similar to Hjortmo et al., we observed double as high folate concentrations in the continuous cultivation compared to the batch flasks experiments, with total folate concentrations of 6.6 mg and 6.7 mg per 100 g dry biomass at day 101 and 133, respectively (c_bm 101, c_bm 133 in Figure 4). The concentrations were comparable to another study in which different strains belonging to *S. cerevisiae* were grown on a glucose-rich medium.[48]

The distribution of the different vitamers was similar to that of batch-cultivation experiments. 5-CH3-H4folate constituted roughly 42-55% of the total folates quantified in the Stage B biomass (Figure 4). H4folate accounted for approximately 19-37% of the total folates. Another dominant form of folate in the samples was 5-CHO-H4folate with 13-16% (Figure 4), and thereby slightly higher in composition than batch-cultivation experiments. 5-CHO-H4folate is known to be a not direct metabolically active vitamer because it has to be converted first, and therefore is assumed to act as a stable folate storage form.[55] It was supposed that excess reduced folates are stored in the form of 5-CHO-H4folate, which can be reactivated when active folate equivalents are needed.^[50b]^ Increased amounts of 5-CHO-H4folate that were produced in our continuous cultivation could, therefore, have resulted from high levels of 5-CH3-H4folate, which is converted as a storage compound. PteGlu was absent in most of the samples. This was expected, as it has no bioactive function in microbes and humans and was rather an artifact from sample preparation, as discussed for *T. kivui*.

H4folate and 5-CH3-H4folate were the most abundant vitamers in the biomass of both *T. kivui* and *S. cerevisiae*. We did not detect folates in the supernatant of the bioreactor (c_sn 101, c_sn 133 in Figure 4). Folates were not released to the medium by *T. kivui,* and we were not able to extract folate from *T. kivui* by cell disruption. Consequently, we can conclude that the yeast in Stage B did not accumulate folates that were produced in Stage A but produced folates itself. The high intracellular folate concentrations in the continuous operation compared to the batch operation emphasize the importance of constantly maintaining *S. cerevisiae* at an optimal growth rate in Stage B. In summary, the high natural inherent abundance of 5-CH3-H4folate in *S. cerevisiae* makes it an ideal choice for consumption, as it supplements the human body with the essential active folate vitamer.

### 3.4. Sustainable processes for food and nutrition can surpass traditional alternatives in terms of nutritional profile

To appraise the marketability and suitability of the described Power-to-Protein system, comparing the yeast product to more widely accepted and standardized food products is fundamental. Therefore, we wanted to compare the nutritional profile of our product to beef, pork, and fish, which are traditional animal-derived protein sources, and lentils, which are plant-based protein sources, because of their widespread consumption worldwide (**Table 4**). The quantities of protein and folate content in the different meat, fish, and vegetable products have been collected from a USDA Database.[56] To account for the uptake of nutrients when each product is consumed in its usual amount, we normalized it to the standard serving size. The standard serving size for meat (beef and pork) is 85 g[57]. We normalized all other products to a serving size of 85 g, and we considered a serving size of 85 g (∼ 6 tablespoons) for nutritional yeast **(Table 4**) to allow a comparison of the protein and folate content to the meat, fish, and vegetable products.

**Table 4.**
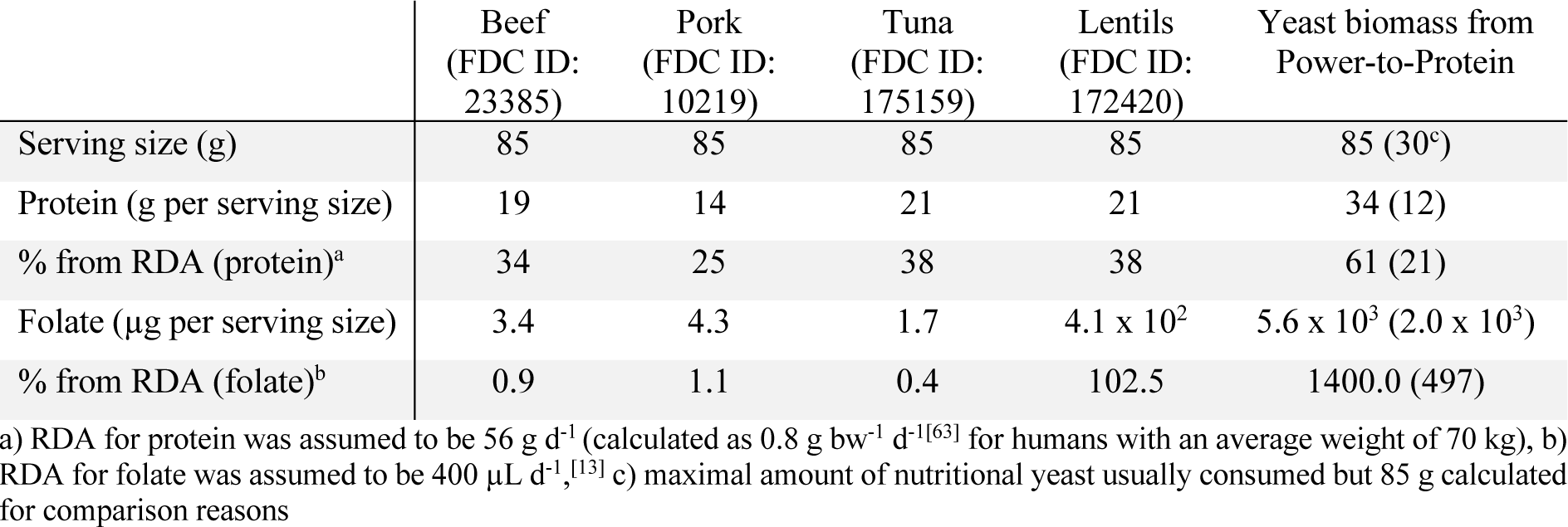
Comparison of the nutritional composition of the yeast biomass from Power-to-Protein to other conventional products by traditional food production systems regarding the recommended daily allowances (RDAs) and the serving size.

85 g of *S. cerevisiae* that was grown in a Power-to-Protein system can meet 61% of the RDA for protein. This value is higher than the same amount of the animal-derived protein products beef, pork, and fish, which provide 34%, 25%, and 38% of the recommended protein values, respectively. As a plant-based product, lentils provide 38% of the recommended daily protein intake. Therefore, based on protein content, we can conclude that yeast biomass is a better protein source than traditional protein sources based on the same serving size. However, high nucleic acid (NA) levels in yeast with about 5-12% of the total biomass composition[58] restrict a high consumption of nutritional yeast by humans as the purines adenine and guanine are metabolized into uric acid, enhancing the risk of gout. A maximum safe amount of 2 g d^-1^ NA is suggested,[59] and therefore a maximum of two tablespoons of flakes, granules, or powder (serving size of ∼30 g, **Table 4** in brackets) of nutritional yeast is often recommended.[60] To mitigate the issues of elevated NA intake with increased serving sizes, yeast biomass can be processed to reduce NA levels. Techniques described to reduce the NA content in SCP are chemical, enzymatic, or thermal treatments, with thermal treatment (72-74°C for 30-45 min) being the most effective.[61] Taking biomass losses of around 30–33% at high-temperature treatments into account,[62] then 41% (instead of 61%) of the daily nutrient requirements for protein (56 g d^-1^ for a 70 kg person[63]) can be met with a serving size of 85 g of yeast from the Power-to-Protein system.

In contrast to being a good source of protein, folate concentrations in animal-derived meat and fish products are low. The folate concentration in the selected meat alternatives at a serving size of 85 g is around ∼1% of the recommended daily allowance (RDA) (**Table 4**). On the contrary, folate content in vegetables, such as lentils, is high, fulfilling 100% of the daily demand for folate. However, the *S. cerevisiae* biomass obtained from Stage B of the Power-to-Protein process with a serving size of 85 g provides 1400% of the total folate RDA. 5-CH3-H4folate is the most common natural folate in food, followed by the formyl forms 5-CHO-H4folate and 10-CHO-H4folate.[64] Due to its bioavailability and bioactivity and no further enzymatic conversion that is needed in the human body, 5-CH3-H4folate was declared as the most desirable vitamer for fortifying food products.^[17a, 65]^ Because 5-CH3-H4folate is the vitamer that was primarily enriched by *S. cerevisiae* (∼50% of the total content), the yeast biomass seems suitable for effective biofortification.

We have yet to determine if the downstream processing to reduce NA levels affects the folate composition in the yeast cells. Literature does report the effects of downstream processing on the stability of the folate vitamers.[66] Folates are reported to be sensitive, especially to heat treatments.[67] For spinach and green bean samples, a 60-80% loss of the initial folate concentrations was observed at 65°C and 85°C and an incubation time of 60 min.[67] While thermal treatment of the yeast biomass may decrease folate concentrations, a serving size of 85 g yeast would still surpass the RDA for folate by far, even taking the 80% loss of folate into account. The effect of downstream processing on the composition of the yeast cells must be further investigated. It would also be possible to extract excess folate and to use it for human consumption elsewhere. In summary, the comparative analysis demonstrates that the intake of 85 g per day of yeast biomass from the Power-to-Protein system can effectively meet the RDAs for both protein and folate. When considering protein provision, yeast biomass stands *on par* or even slightly better with conventional animal-and plant-based sources while surpassing them in terms of folate content. This underscores the potential of the Power-to-Protein system as a valuable and nutritionally competitive alternative for meeting essential dietary requirements.

## 4. Conclusion

Leveraging yeast biomass as a protein alternative has the potential to alleviate a range of global challenges, including environmental degradation, food security, and health. It is essential to engage developing nations in the creation of upcoming technologies aimed at reducing carbon emissions in global food production. Such nations, experiencing rapid population growth, urgently require an alternative food production system, such as Power-to-Protein, to overcome food scarcity and reduce the pressure on current food production systems. Yeast protein, while delivering essential vitamins, diminishes the risk of malnutrition. Our investigations have encouraged the development of the Power-to-Vitamin system to ensure two objectives: **(1)** protein supply for nutrition security; and **(2)** adequate vitamin intake. We showed that folate production from renewable electric power and CO2 was possible for both our anaerobic stage (*T. kivui*) and our aerobic stage (*S. cerevisiae*). Even though there is no evidence of microbe-mediated biofortification of folate into yeast through *T. kivui* in Stage A, *S. cerevisiae* was itself capable of folate bioproduction while using acetate as a carbon source. Nevertheless, a more in-depth investigation is necessary to thoroughly explore how to leverage the high folate concentrations in *T. kivui* either by adding cells (if food regulations allow) or extracted folate to human food.

We observed that folate production depended on the growth rate for *S. cerevisiae*, with a continuous operating mode optimal to achieve high total folate concentrations. This corroborates previous studies, but we used acetate as the sole carbon source here. Nevertheless, optimizations of the Power-to-Protein system are needed to enhance biomass productivity and secure steady-state production over a long period. By this, unwanted alterations in the growth phase of the yeast in Stage B are prevented, thereby, stabilizing the production of folates. Furthermore, an investigation on medium composition is needed because we should only procure specific necessary vitamins to enhance sustainability further and reduce the costs of the overall process. A first simple economic model projected the Power-to-Protein system to feed 10 billion people with ten thousand fermenters with a capacity of 3100 m^3^[8] after optimizing the protein productivity to 1.25 g L^-1^ h^-1^. This motivates the continuation of research to further upgrade it into a Power-to-Vitamin process that focuses on overall nutrient administration in addition to providing protein.

## Acknowledgments

The authors acknowledge support from the Alexander von Humboldt Foundation in the framework of the Alexander von Humboldt Professorship that is endowed by the Federal Ministry of Education and Research in Germany and the CO2 Research Center funded by the Novo Nordisk Foundation with grant number NNF21SA0072700 (both to L. T. A.). In addition, the vitamin analyses were financed by a validation project of SPRIND GmbH, contract number 027/2022, and additional support came from the Federal Ministry of Education and Research with grant number 031B0857D. The authors thank Lucas Reiner for help with the folate analysis.

## TOC

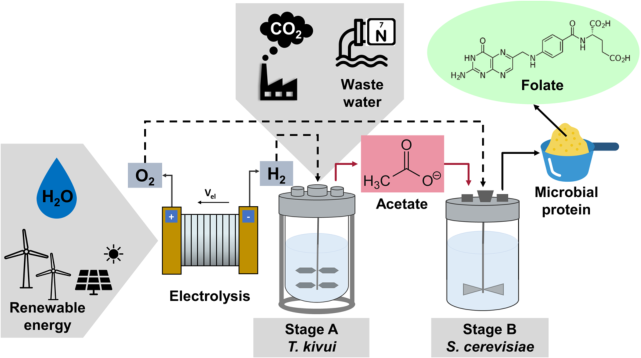

Microbial protein from renewable power and CO2 as an alternative food source has the potential to address major challenges that are associated with conventional agriculture. This study showed that yeast enriches folate when growing on biologically produced acetate in a two-stage bioreactor system. Producing protein and vitamins promises to overcome food scarcity and micronutrient deficiency (icons taken from Flaticon).

A. Lisa Marie Schmitz, A. Nicolai Kreitli, B. Lisa Obermaier, B. Nadine Weber, B. Michael Rychlik, A.C.D.E.F. Largus T. Angenent*

## Supporting Information

Supporting information is available from the Wiley Online Library or from the author.

