## Supporting information for "Power-to-Vitamins: Producing Folate (Vitamin B_9_) from Renewable Electric Power and CO_2_ with a Microbial Protein System"

### Experimental Procedures

**MS medium:**

<u>MS mineral salts (10x):</u> NaCl (4.5 g $L^{-1}$), $K_2HPO_4$ (1.7 g $L^{-1}$), $NH_4Cl$ (1.9 g $L^{-1}$), MgCl x $6H_2O$ (0.8 g $L^{-1}$), $CaCl_2$ (0.6 g $L^{-1}$), and $KH_2PO_4$ (2.3 g $L^{-1}$)

<u>Trace element solution MS (1000x):</u> $MgSO_4$ x 7 $H_2O$ (30 g $L^{-1}$), $MnSO_4$ x $H_2O$ (5 g $L^{-1}$), NaCl (10 g $L^{-1}$), $FeSO_4$ x 7 $H_2O$ (1 g $L^{-1}$), $CoCl_2$ x 6 $H_2O$ (1.8 g $L^{-1}$), $CaCl_2$ x 2 $H_2O$ (1 g $L^{-1}$), $ZnSO_4$ x 7 $H_2O$ (1.8 g $L^{-1}$), $CuSO_4$ x 5 $H_2O$ (0.1 g $L^{-1}$), KAl $(SO_4)_2$ x 12 $H_2O$ (0.18 g $L^{-1}$), $H_3BO_3$ (0.1 g $L^{-1}$), $Na_2MoO_4$ x 2 $H_2O$ (0.1 g $L^{-1}$), $(NH_4)_2Ni(SO_4)_2$ x 6 $H_2O$ (2.8 g $L^{-1}$), $Na_2WO_4$ x 2 $H_2O$ (0.1 g $L^{-1}$), and $Na_2SeO_4$ (0.1 g $L^{-1}$)

**Yeast nitrogenous base medium (YNB):**

<u>YNB mineral salts:</u> $(NH_4)_2SO_4$ (5 g $L^{-1}$), $KH_2PO_4$ (1 g $L^{-1}$), $MgSO_4$ x $7H_2O$ (1.025 g $L^{-1}$), NaCl (0.1 g $L^{-1}$), and $CaCl_2$ (0.132 g $L^{-1}$)

<u>YNB trace element solution (100x):</u> $H_3BO_3$ (50 mg $L^{-1}$), $CuSO_4$ x $5H_2O$ (6.25 mg $L^{-1}$), KI (10 mg $L^{-1}$), $FeCl_3$ x 6 $H_2O$ (33.33 mg $L^{-1}$), $MnSO_4$ (44.77 mg $L^{-1}$), $Na_2MoO_4$ x $2H_2O$ (23.51 mg $L^{-1}$), and $ZnSO_4$ x $7H_2O$ (71.3 mg $L^{-1}$)

<u>YNB vitamin solution (100x):</u> myo-inositol (200 mg $L^{-1}$), thiamine HCl (32 mg $L^{-1}$), pyridoxine HCl (40 mg $L^{-1}$), D-pantothenic acid hemicalcium salt (40 mg $L^{-1}$), biotin (0.2 mg $L^{-1}$), riboflavin (20 mg $L^{-1}$), aminobezoic acid (20 mg $L^{-1}$), and nicotinic acid (40 mg $L^{-1}$)

**Modified 2xP7-medium:**

<u>P7 mineral salts (33.33x):</u> NaCl (80 g $L^{-1}$), $NH_4Cl$ (100 g $L^{-1}$), KCl (10 g $L^{-1}$), $KH_2PO_4$ (10 g $L^{-1}$), MgCl x 6 $H_2O$ (33.7 g $L^{-1}$), and $CaCl_2$ x 2 $H_2O$ (4 g $L^{-1}$)

P7 trace element solution (100x): nitrilotriacetic acid (NTA) (2 g $L^{-1}$), $MnCl_2$ x $4H_2O$ (1.32 g $L^{-1}$), $(NH_4)_2Fe(SO_4)_2$ (0.8 g $L^{-1}$), $CoCl_2$ x $6H_2O$ (0.2 g $L^{-1}$), $ZnSO_4$ x $7H_2O$ (0.356 g $L^{-1}$), $CuCl_2$ x $2H_2O$ (20 mg $L^{-1}$), $NiCl_2$ x $6H_2O$ (20 mg $L^{-1}$), $Na_2MoO_4$ (20 mg $L^{-1}$), $Na_2SeO_3$ x $5H_2O$ (27.74 mg $L^{-1}$), and $Na_2WO_4$ x $2H_2O$ (22 mg $L^{-1}$)

P7 vitamin solution (100x): pyridoxine HCl (10 mg $L^{-1}$), riboflavin (5 mg $L^{-1}$), D-pantothenic acid hemicalcium salt (5 mg $L^{-1}$), aminobezoic acid (5 mg $L^{-1}$), nicotinic acid (5 mg $L^{-1}$), folic acid (2 mg $L^{-1}$), mercaptoethansulfonic acid (10 mg $L^{-1}$), thiamine HCl (5 mg $L^{-1}$), DL-6-8-ditihiooctan acid (5 mg $L^{-1}$), cobalamin (5 mg $L^{-1}$), and biotin (2 mg $L^{-1}$)

**Setting up the Stage A bioreactor**

The Stage A bioreactor vessel was autoclaved containing MS mineral salts and trace elements (1x concentrated). Once, the vessel cooled down to room temperature, the vessel was first connected to an air gas line for oxygen sensor calibration and then to the $H_2/CO_2$ gas line overnight to ensure anaerobic conditions. The next day, the vitamin and reducing agent were added to the Stage A bioreactor. MS medium (1 x concentrated) contained 10 % (vol/vol) MS, mineral salts (10 x), 0.1% (vol/vol) trace element solution, L-cysteine (0.5 g $L^{-1}$), 0.1% (vol/vol) $(NH_4)2Ni(SO_4)_2$ (0.2%, w/v), 0.1% (vol/vol) $FeCl_2$ x 4 $H_2O$ (0.2%, w/v), and was adjusted to a pH of 6.4. Later, the bioreactor was inoculated with *T. kivui* pre-culture to reach an initial $OD_{600}$ of 0.01. The continuous mode of operation was initiated on day 4 with a cell-recycling module. The module was sterilized by autoclaving at 121 °C, 2 bar for 50 min, and rinsed thoroughly with sterile, anoxic distilled water.

**Setting up the Stage B bioreactor**

The Stage B bioreactor vessel was sterilized by autoclaving YNB mineral salts and trace elements (both 2-times concentrated) filled up with $dH_2O$ to a final volume of 750 mL. Autoclaved medium was connected to a gas line providing compressed air. The vitamin solution was added once the Stage B bioreactor had cooled down to room temperature. The reactor was filled up with effluent from Stage 1 to a final volume of 1,500 mL. 1-times concentrated YNB medium contained YNB mineral salts, 1 % (vol/vol) YNB trace element solution (100 x), and 1% (vol/vol) YNB vitamin solution (100x). Because a continuous pH control was provided, no MES-buffer was added for bioreactor cultivations. The medium was inoculated with *S. cerevisiae* pre-culture grown in YPD medium to a start $OD_{600}$ of 0.05. Once the culture reached an exponential phase of growth, a continuous mode of operation was started. Filtered effluent from Stage 1 was pumped to the Stage B reactor. A by-pass medium feed line containing 2- or 4-times concentrated YNB medium was added to the Stage B bioreactor. The ratio of effluent to medium varied during the different Periods. The tubes of all multichannel pumps were sterilized by autoclaving.

**Cultivation conditions *Clostridium ljungdahlii***

Cultures of *C. ljungdahlii* PETC (DSM 13528) were first grown from frozen cryo-cultures in 100 mL Reinforced Clostridial Medium (RCM) medium with $N_2$ atmosphere at 37°C without shaking for 24 h. RCM contained fructose (5 g $L^{-1}$), yeast extract (3 g $L^{-1}$), beef extract (10 g $L^{-1}$), peptone (10 g $L^{-1}$), NaCl (5 g $L^{-1}$), soluble starch (1 g $L^{-1}$), sodium acetate (3 g $L^{-1}$), resazurin (0.025 %), cysteine (0.5 g $L^{-1}$). 1 mL of cultures was transferred to 100-mL injection bottles containing 50 mL modified 2xP7 medium with yeast extract (0.5 g $L^{-1}$), MES (0.04 g $L^{-1}$), fructose (0.1 g $L^{-1}$), pH of 5.5 and $N_2$ atmosphere. Bottles were cultivated at 37°C and without shaking for 72 h. Stage A bioreactor was operated in batch mode containing 1-times concentrated 2xP7 medium with yeast extract (1 g $L^{-1}$). 2-times concentrated 2xP7-medium was fed in continuous operating mode with a feeding rate of ~22 mL $h^{-1}$, corresponding to a dilution rate of 0.011 $h^{-1}$. In Period II the 2xP7 medium contained yeast extract (0.5 g $L^{-1}$) and complete vitamin addition. In Period III the 2xP7 medium contained no yeast extract and folic acid and cobalamin were omitted from the medium. A cell dry weight (CDW) correlation factor of 242 mg CDW $OD_{600}^{-1}$ $L^{-1}$ for *C. ljungdahlii* was used.

**Analytical methods**

A gas chromatography system SRI 8610C was used to analyze the gas compositions (hydrogen and carbon dioxide). Samples were taken with a 500-µL gastight syringe (Hamilton, Reno, USA) and a HaySep-D packed Teflon column (length 3 m, outer diameter 1/8") (SRI GC, Torrance, USA) at 25°C and nitrogen as carrier gas were used. The instrument was equipped with a thermal coupled detector (TCD) and a flame ionization detector (FID).

The protein concentrations were analyzed using the BCA protein assay (Pierce™ BCA Protein Assay Kit, Thermo Fisher Scientific™, Waltham, MA, USA) according to the manual and by protein determination according to Lowry [33]. The Lowry method combines the reactions of cupric ions with the peptide bonds under alkaline conditions (the Biuret test) with the oxidation of aromatic protein residues. Samples and BSA standards (0 µg mL$^{-1}$, 112.5 µg mL$^{-1}$, 225 µg mL$^{-1}$, 450 µg mL$^{-1}$, 675 µg mL$^{-1}$, 900 µg mL$^{-1}$, 1125 µg mL$^{-1}$, 1350 µg mL$^{-1}$) were boiled in a heating block at 100°C for 10 min and cooled on ice. Samples were diluted if needed. 10 µl of each BSA standard or sample was transferred into a 96-well plate. 200 µl of reagent A ($Na_2CO_3$ (0.2 M), NaOH (0.1 M), $CuSO_4$ (0.6 mM), and NaK-tartrate (0.7 mM)) was added to each well. Blank values at an absorbance of 750 nm were read before adding 20 µL of reagent B (1 M Folin-Ciocalteu's phenol reagent) to every well. The plate was incubated in the dark at room temperature for 15 minutes and measured again at 750 nm. Concentrations were calculated by subtracting the background values and correlating them with the standard curve.

Analysis of ammonium concentrations in the medium and effluent of Stage A was performed by the Nessler method using an Ammonia High Range Reagent Kit (Hanna Instruments, Smithfield, RI, USA). Samples were diluted 1:20 with $dH_2O$ prior to measurement. Trace elements in the medium and effluent of Stage A were analyzed by inductively coupled plasma mass spectrometry (ICP-MS) on an Agilent 7900 ion-coupled plasma mass spectrometer instrument (Agilent, Santa Clara, USA). The samples were diluted 1:100 in 1% HNO3, which was also the matrix. $^{24}$Mg, $^{48}$Ca, $^{55}$Mn, $^{56}$Fe, $^{59}$Co, $^{60}$Ni, $^{66}$Zn, $^{78}$Se, $^{95}$Mo were quantified. Helium was used as a carrier gas. Data analysis was performed using the MassHunter software (Agilent, Santa Clara, USA).

Freeze-drying of the biomass samples for vitamin analyses was performed with a double-chamber Alpha 1-4 LSCplus laboratory freeze-dryer (Martin Christ Gefriertrocknungs-anlagen GmbH, Osterode am Harz, Germany). Frozen samples were placed in the drying chamber above the -50°C ice condenser chamber, with both chambers maintained at a pressure of 0.1 mbar. Samples were freeze-dried until the pressure within the chambers no longer increased and remained stable at 0.1 mbar.

### Supplementary data

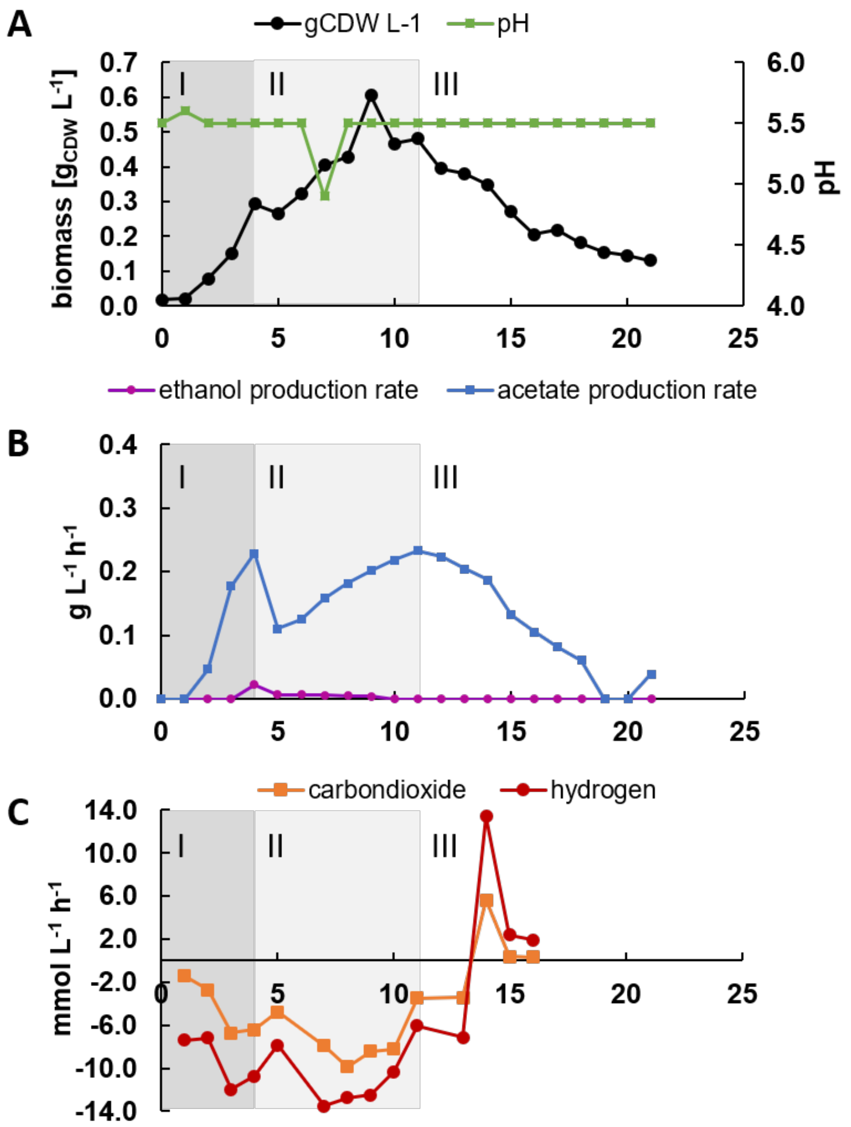


**Figure S1.** Performance of Stage 1 with *C. ljungdahlii*. (A) Biomass in grams cellular dry weight (CDW) per liter (black) and pH (green); (B) Production rate of acetate in g $L^{-1}$ $h^{-1}$ (blue); (C) Consumption rates for $H_2$ (red) and $CO_2$ (orange) in mmol $L^{-1}$ $h^{-1}$. Period I: batch operation with 2xP7 medium containing 1 g $L^{-1}$ yeast extract and complete vitamin addition, period II: continuous operation with 2xP7 containing 0.5 g $L^{-1}$ yeast extract and complete vitamin addition, period III: continuous operation without yeast extract and vitamin solution without folate and cobalamin.

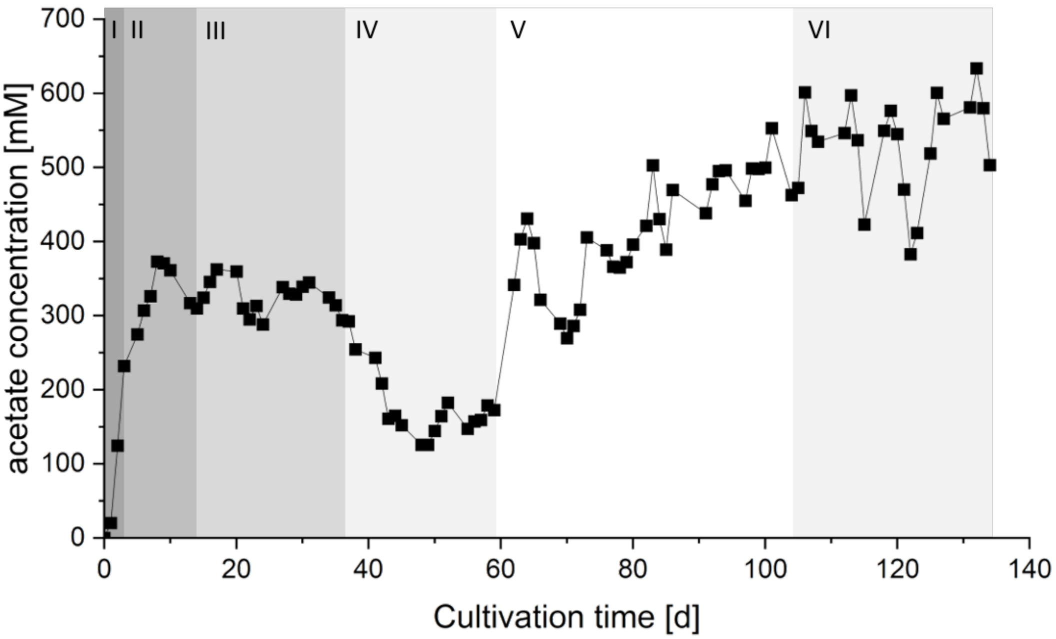


**Figure S2.** Acetate concentrations in Stage A in mM during the cultivation time.

**Table S1.** Trace element analysis by ICP-MS and ammonium analysis by the Nessler method at Day 123 and Day 120, respectively.

| Compound analyzed | MS Medium [4x] | Stage A effluent |
|---|---|---|
| $NH_4^+$ | 760 mg $L^{-1}$ | 172 mg $L^{-1}$ |
| $^{24}Mg$ | 51.8 x $10^3$ ppb | 41.2 x $10^3$ ppb |
| $^{48}Ca$ | 92.0 x $10^3$ ppb | 75.2 x $10^3$ ppb |
| $^{55}Mn$ | 64.3 x $10^2$ ppb | 46.4 x $10^2$ ppb |
| $^{56}Fe$ | <LoQ | <LoQ |
| $^{59}Co$ | 20.1 x$10^2$ ppb | 14.8 x $10^2$ ppb |
| $^{60}Ni$ | 31.8 x $10^2$ ppb | 25.2 x $10^2$ ppb |
| $^{66}Zn$ | 16.2 x $10^2$ ppb | 47.2 x $10^1$ ppb |
| $^{78}Se$ | 36.1 x $10^1$ ppb | 33.8 x $10^1$ ppb |
| $^{95}Mo$ | 13.5 x $10^1$ ppb | 74.4 ppb |

**Table S2.** Details of all the folate vitamers measured within each sample. Folates were extracted from the samples in duplicates. Values are means of duplicates with standard deviation provided for each vitamer in each sample. LoQ = limit of quantification, n.d. = not detected.

| **Name** | **PteGlu** [µg/100g] ± RSD [%] | **$H_4$folate** [µg/100g] ± RSD [%] | **5-$CH_3$-$H_4$folate** [µg/100g] ± RSD [%] | **5-CHO-$H_4$folate** [µg/100g] ± RSD [%] | **10-CHO-PteGlu** [µg/100g] ± RSD [%] | **Total folate content [µg/100g] ± RSD [%]** |
|---|---|---|---|---|---|---|
| Sce b_f1 | $1.0 \times 10^1$ ± 6.4 | $1.2 \times 10^3$ ± 6.4 | $1.6 \times 10^3$ ± 6.2 | $2.5 \times 10^2$ ± 4.1 | $6.4 \times 10^1$ ± 4.0 | $3.2 \times 10^3$ ± 6.0 |
| Sce b_f2 | 7.7 ± 8.3 | $1.2 \times 10^3$ ± $1.1 \times 10^1$ | $1.8 \times 10^3$ ± 3.9 | $3.3 \times 10^2$ ± 6.4 | $6.3 \times 10^1$ ± 5.1 | $3.5 \times 10^3$ ± 1.4 |
| Tki c_bm | $3.5 \times 10^3$ ± 4.1 | $3.2 \times 10^3$ ± 4.6 | $3.5 \times 10^3$ ± 8.3 | $2.4 \times 10^3$ ± $1.0 \times 10^1$ | $2.0 \times 10^2$ ± 4.6 | $1.3 \times 10^4$ ± 2.1 |
| Tki c_sn | < LoQ | $2.9 \times 10^1$ ± 3.8 | < LoQ | 2.4 ± 5.5 | < LoQ | $3.2 \times 10^1$ ± 3.7 |
| Sce c_bm 101 | $4.4 \times 10^2$ ± 1.9 | $1.3 \times 10^3$ ± 0.8 | $3.6 \times 10^3$ ± 1.2 | $8.3 \times 10^2$ ± 0.6 | $4.3 \times 10^2$ ± 3.8 | $6.6 \times 10^3$ ± 0.1 |
| Sce c_sn 101 | < LoQ | n.d. | < LoQ | < LoQ | < LoQ | < LoQ |
| Sce c_bm 133 | $8.8 \times 10^1$ ± 3.7 | $2.5 \times 10^3$ ± 3.1 | $2.8 \times 10^3$ ± 1.7 | $1.1 \times 10^3$ ± 0.1 | $2.8 \times 10^2$ ± 2.3 | $6.7 \times 10^3$ ± 2.0 |
| Sce c_sn 133 | < LoQ | n.d. | < LoQ | < LoQ | < LoQ | < LoQ |

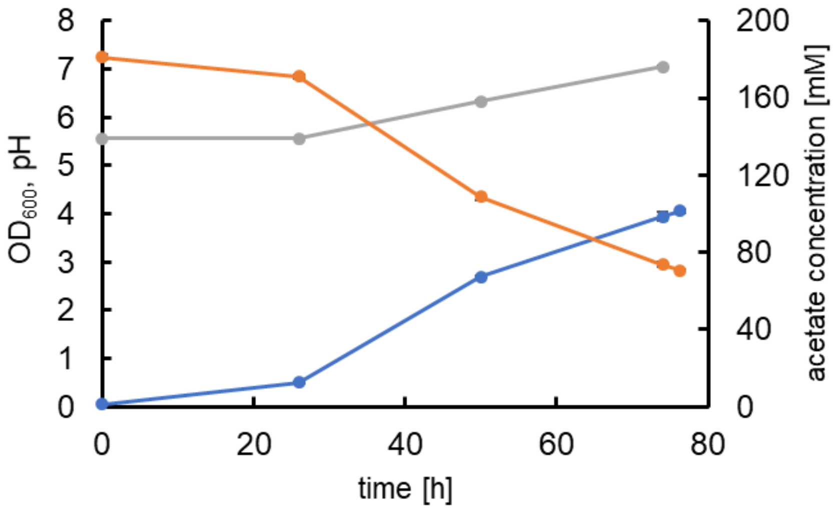


**Figure S3.** Growth of *S. cerevisiae* in batch cultivation experiments on acetate as sole carbon source. YNB medium without folate contained 200 mM acetate. $OD_{600}$ (blue), acetate concentration in mM (orange), pH (grey). Values are the mean of triplicated with SD. Cells were harvested after 76 h of cultivation to determine folate concentrations.
